# Group B *Streptococcus* drives major transcriptomic changes in the colonic epithelium

**DOI:** 10.1101/2023.01.24.525471

**Authors:** Kristen Domínguez, April K. Lindon, Justin Gibbons, Sophie E. Darch, Tara M. Randis

## Abstract

Group B Streptococcus (GBS) is a leading cause of infant sepsis worldwide. Colonization of the gastrointestinal tract is a critical precursor to late-onset disease in exposed newborns. Neonatal susceptibility to GBS intestinal translocation stems from intestinal immaturity; however, the mechanisms by which GBS exploits the immature host remain unclear. β-hemolysin/cytolysin (βH/C) is a highly conserved toxin produced by GBS capable of disrupting epithelial barriers. However, its role in the pathogenesis of late-onset GBS disease is unknown. Our aim was to determine the contribution of βH/C to intestinal colonization and translocation to extraintestinal tissues. Using our established mouse model of late-onset GBS disease, we exposed animals to GBS COH-1 (WT), a βH/C-deficient mutant (KO), or vehicle control (PBS) via gavage. Blood, spleen, brain, and intestines were harvested 4 days post-exposure for determination of bacterial burden and isolation of intestinal epithelial cells. RNA-sequencing was used to examine the transcriptomes of host cells followed by gene ontology enrichment and KEGG pathway analysis. A separate cohort of animals was followed longitudinally to compare colonization kinetics and mortality between WT and KO groups. We demonstrate that dissemination to extraintestinal tissues occurred only in the WT exposed animals. We observed major transcriptomic changes in the colons of colonized animals, but not in the small intestines. We noted differential expression of genes that indicated the role of βH/C in altering epithelial barrier structure and immune response signaling. Overall, our results demonstrate an important role of βH/C in the pathogenesis of late-onset GBS disease.

## Introduction

Group B *Streptococcus* (*Streptococcus agalactiae*; GBS) is a leading cause of neonatal sepsis worldwide. Although there has been a significant decline in the incidence of early-onset sepsis in countries that have implemented maternal GBS screening and intrapartum antibiotic prophylaxis, this strategy has not affected the incidence of late-onset sepsis. In fact, late-onset sepsis comprises a significant portion of invasive GBS disease in many regions of the world(1) and remains a leading cause of neonatal meningitis(2).

GBS is a pathobiont of the intestinal tract, able to asymptomatically colonize the intestines in exposed newborns, but also capable of translocating through intestinal barriers via the paracellular(3) and transcellular(4) routes leading to invasive disease. Neonates are particularly susceptible to GBS invasive disease. Animal studies demonstrate intestinal immaturity contributes to this susceptibility. This includes incomplete intestinal epithelial junction polarization and an immature intestinal microbiome leading to increased GBS colonization, dissemination, and meningitis(5). However, most infants exposed to GBS do not succumb to invasive disease. The mechanisms by which GBS exploits the immature intestinal barriers are poorly understood and hinder the development of strategies to prevent disease in more than 300,000 infants globally(6).

β-hemolysin/cytolysin (βH/C) is a highly conserved, pore-forming, rhamnolipid toxin produced by GBS that contributes significantly to its pathogenicity. Rhamnolipid toxins produced by other pathogens are known to enhance bacterial paracellular translocation via disruption of epithelial barriers(7). Indeed, βH/C plays a critical role in the pathogenesis of pneumonia, meningitis, urinary tract infections, and chorioamnionitis by disrupting epithelial barriers and inducing host inflammation(8–14); however, its role in disrupting the intestinal epithelium, remains unexplored.

In this study, our objective was to explore GBS-host interactions in the intestinal environment and determine the contribution of the βH/C toxin. We hypothesized that the βH/C toxin directly modulates gene expression in intestinal epithelial cells and contributes to subsequent translocation of intestinal barriers. Using our robust murine model for late-onset GBS disease(15) we demonstrate that GBS colonization induces differential gene expression primarily in the colonic epithelium, with toxin-specific effects on genes regulating epithelial cell structure and immune signaling.

## Methods

### GBS strains and growth

GBS COH-1 (serotype III, ST-17) (WT) and its isogenic βH/C deficient mutant, COH1delcylE (KO, gifted from Adam J. Ratner, MD), were incubated in liquid tryptic soy broth at 37°C overnight. The COH-1 strain of GBS (serotype III, ST-17) was used for all experiments. Bacteria were grown to stationary phase at 37°C in Trypticase soy (TS) broth and enumerated by plating on TS agar or CHROMagar^™^ StrepB plates. Cultures were centrifuged and pellet resuspended in sterile Dulbecco’s phosphate-buffered saline (PBS) for immediate use in mouse infections.

### Mouse model

All experiments were performed in accordance with the University of South Florida Institutional Animal Care and Use Committee (IACUC). A postnatal GBS colonization model was used as previously described(15) with few modifications. Briefly, adult (8 to 10 weeks old) C57BL6/J mice were purchased from Jackson Laboratories. After acclimating, mice were mated and liveborn offspring were assessed daily. Bacterial cultures were grown overnight to the stationary phase, centrifuged, and resuspended in sterile PBS to a final concentration of 10^8^-10^9^ CFU/ml. On day of life 10±1, mice were infected via oral gavage with either WT or KO GBS resuspended in 100 μl of PBS using a sterile feeding tube. A sham-infected cohort was inoculated with 100 μl of vehicle control (PBS). All littermates were in the same study group and pups remained with their biologic, noncolonized dam for the remainder of the experiment. To obtain long-term infection data of mortality and colonization, we monitored, weighed, and collected rectal swabs from mice every 2-3 days. Swabs were stored in PBS and plated on GBS selective media (described above), which were incubated overnight at 37°C.

### Tissue processing for GBS detection and host cell gene expression

In a separate cohort, mice were euthanized, and blood, spleen, brain, small intestine, and colon tissues were extracted to determine GBS burden 4 days post-gavage. This timepoint was selected to reflect sustained colonization rather than immediate exposure to GBS and is typically the point at which we observe the greatest mortality. Tissues were placed in PBS and gently homogenized using the Bullet Blender^®^ (Next Advance inc.) for 30 seconds on speed 3. Serial dilutions of tissue homogenates were plated on selective media (CHROMagar ^™^ StrepB) to determine GBS CFU’s in tissue and blood samples (100μL) were streaked onto selective media to determine presence of GBS. A separate cohort of each of the 3 study groups (PBS, WT, KO) was euthanized 4 days post-gavage. Whole small intestine and colon samples were collected and processed for intestinal epithelial cell isolation using an established protocol(16). Briefly, RNA was extracted with the QIAGEN RNeasy^®^ Kit and then purified with the Invitrogen TURBO^®^ DNA-free kit. Purified RNA samples were shipped to Novogene Co., Ltd. for RNA sequencing (via the Illumina platform) and library creation. A subset of RNA samples was retained to perform targeted qRT PCR to confirm select differentially regulated genes from RNA-seq analysis using TaqMan^™^ Gene Expression Assays (Supplemental Table I).

### RNA-seq data analysis

The transcriptome similarity between all samples was measured using PCA plots of the top 500 most variable genes after the variance stabilizing transformation from DESeq2 Version 1.34.0 was applied(17). Genes differentially expressed between each comparison group were found using DESeq2 Version 1.34.0 using default settings. Genes with a false discovery rate (FDR) less than or equal to 0.1 were considered differentially expressed. The differentially expressed genes were assessed for gene ontology enrichment. The gene ontology annotations were obtained from the AnnotationDBI and org.Mm.eg.db R packages (version 1.56.2 and 3.14.0, respectively)(18, 19). The R package topGO version 2.46.0 was used to calculate gene ontology enrichment using the weight01 algorithm and the Fisher’s exact test(20). Gene ontology categories with a p-value of less than or equal to 0.05 were considered enriched.

Gene set enrichment analysis was also performed on specific sets of genes obtained from a previous study on the effects of microbiota on the infant gut(21) and selected KEGG pathways (signal transduction, signaling molecules and interaction, transport and catabolism, cell growth and death, cellular community – eukaryotes, cell motility, immune system, and infectious disease: bacterial). The gene ontology gene sets genes were obtained using the AnnotationDBI and org.Mm.eg.db R packages (version 1.56.2 and 3.14.0, respectively)(18, 19). The KEGG gene sets were obtained from the R package gage (version 2.44.0) using the kegg.gsets function(22). Genes that had a median expression level greater than the overall genome wide median expression level were used in this analysis. For this analysis, the gene expression count data was normalized using DESeq2 normalization(17) and the KStest function from the R package GSAR (version 1.28.0) was used to test for changes in the gene set mean expression(23).

### Statistical analysis

Statistics were calculated using GraphPad Prism 9 software. Survival and colonization duration data were compared using Kaplan-Meier curve analysis with the Log-Rank test. Dissemination data was analyzed using Fisher’s exact test. Tissue burdens (CFU/gram tissue) were compared using nonparametric Mann-Whitney test. We analyzed qRT-PCR data using the ΔΔ-CT method and One-way ANOVA with uncorrected Fisher’s LSD multiple comparisons test. The function KStest from the R package GSAR version 1.28.0 was used to assess changes in expression within the gene sets between each of the samples(23). Expression differences of pathways with an FDR of less than or equal to 0.1 were considered to be differentially expressed.

### Data Availability

The raw RNA-seq data and the DESeq2 processed data have been deposited in the GEO database.

## Results

### GBS induces major transcriptomic changes primarily in the colon, with toxin-specific effects

To gain an in-depth view of GBS-host interactions in the intestinal epithelium, we used RNA-seq to examine host gene expression 4 days post-exposure. We determined the gene expression profiles of the intestinal epithelial cells from both the colon and small intestine in mice exposed to PBS, WT, or KO. Of note, we demonstrate no significant differences in bacterial burden in the intestine of either of GBS exposed groups (WT or KO) (Figure 1A) yet, we observed striking differences in host gene expression profiles. Extraintestinal dissemination (recovery of GBS from blood, spleen or brain) at this timepoint occurred exclusively in the WT-exposed group (9.8%), though this difference did not reach statistical significance (P= 0.14, Fisher’s exact test) (Figure 1B). In the colon, we noted 6,822 and 3,364 significant differentially expressed genes in the WT vs PBS and WT vs KO groups, respectively, with 2,699 genes overlapping. There were no significant differentially expressed genes in the KO vs PBS colon comparison. We found little differences among all groups in the epithelial cells isolated from the small intestines (Figure 1C). There were much greater differences in gene expression between PBS and WT, with changes in the KO group that indicate toxin-specific functions (Figure 1D).

**Figure 1.**
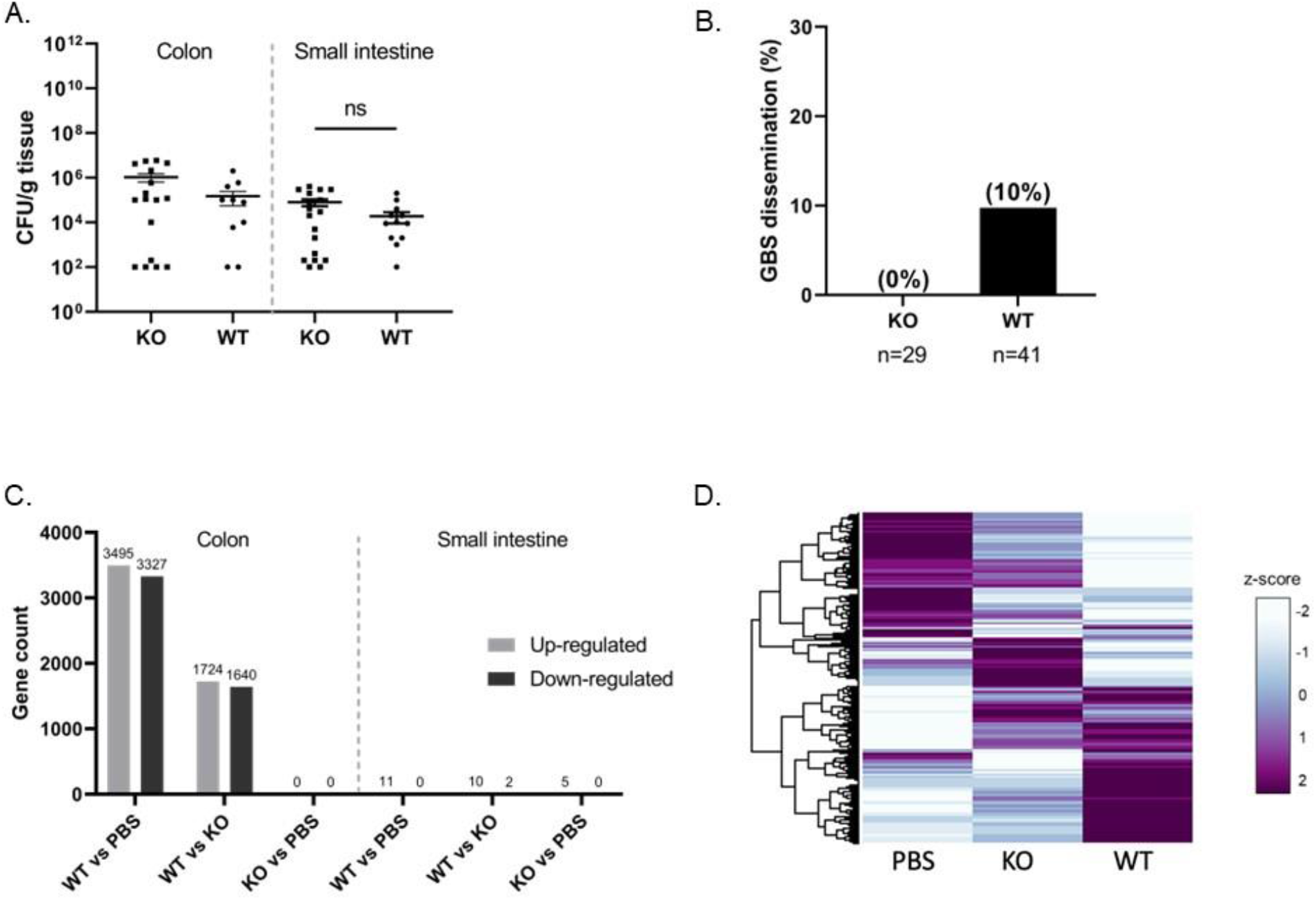
GBS induces major transcriptomic changes primarily in the colon, with toxin-specific effects. A) GBS intestinal burden in the colon and small intestine by group (WT or KO) displayed as CFU/g tissue. B) Percent of mice in either WT or KO where GBS was isolated from extra intestinal tissues (spleen or brain) and/or blood. (n=41 WT, n=29 KO). C) Gene counts of the significant up and down regulated genes in the WT vs PBS, WT vs KO, and KO vs PBS comparisons in both the colon and small intestine. D) Heatmap displaying the normalized z-scores of the gene expression from PBS, KO, and WT exposed mice colons. (Small intestine: n=4 WT, n=4 KO, n=4 PBS; Colon: n=5 WT, n=5 KO, n=4 PBS).

Among the significantly differentially expressed genes in the colon, we identified the top 10 genes with the greatest fold change (Figures 2A-B). Many of these genes have been previously reported to regulate intestinal homeostasis and barrier function suggesting important functional changes caused by GBS colonization.

**Figure 2.**
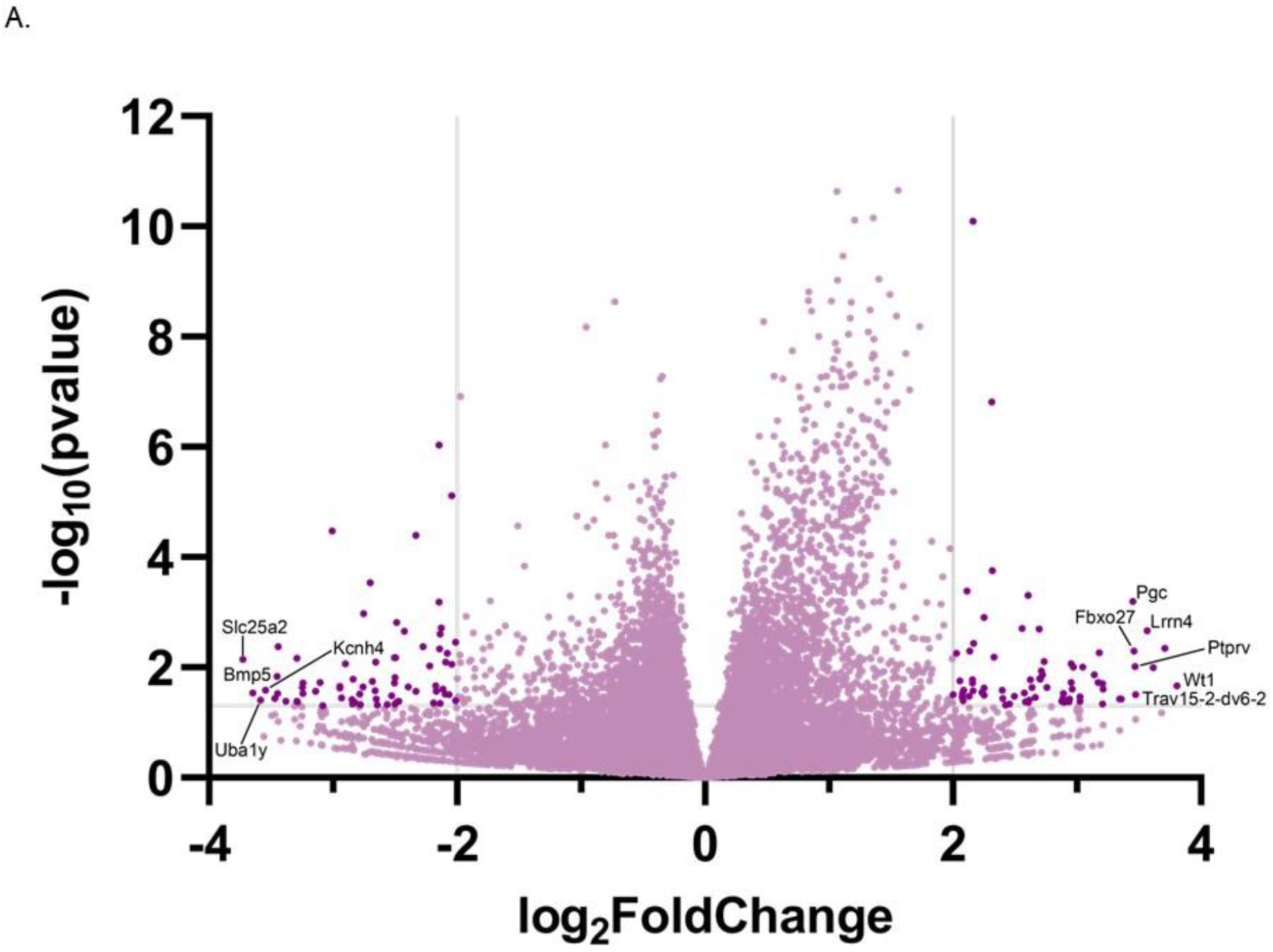
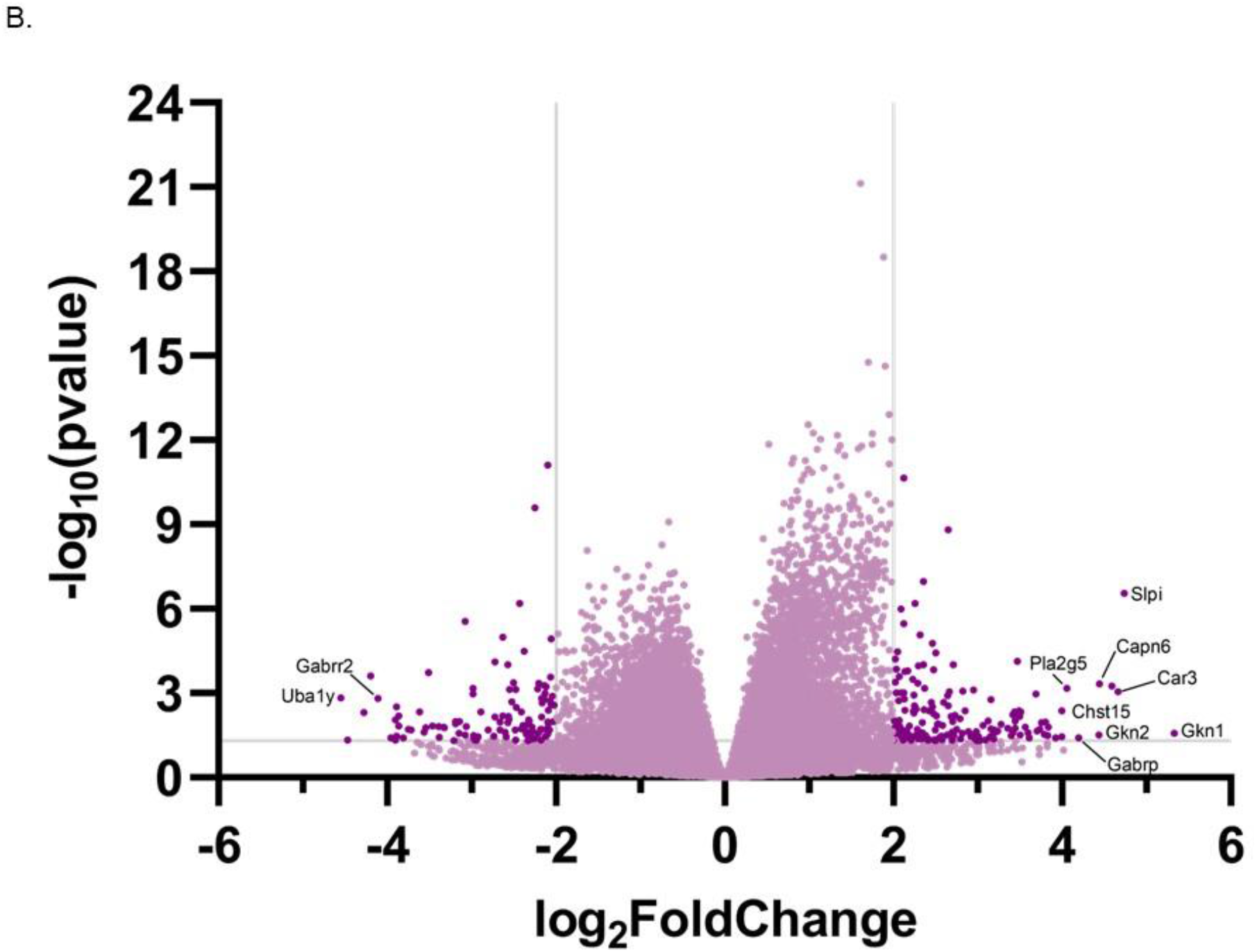
Differential gene expression among PBS, WT, and KO exposed mice in the colon. Volcano plots displaying the up and down regulated genes in the C) WT vs PBS and D) WT vs KO comparisons. The horizontal gray line denotes P value = 0.05; the points above this line are significant. Vertical gray lines denote log2foldchange of −2 or 2; the points less than or greater than those values, respectively, have more substantial changes in their gene expression. Dark purple points indicate those genes that are within these parameters. Light purple points indicate genes that fall outside of these parameters. The top 10 differentially regulated genes, based on absolute log_2_foldchange, are labeled on the graph.

### Transcriptomic changes are associated with epithelial barrier function and immune regulation

To analyze functional changes of the differentially expressed genes, we used GO enrichment analysis. Here, we found significant changes in molecular processes related to transcription, translation, and immune regulation in both the WT vs PBS (Figure 3A) and WT vs KO comparisons (Figure 3B). When looking at the top 10 drivers of gene expression (see Figures 2A-B), we also noted GO terms related to antibacterial responses (*Slpi*, *Car3*, *Gkn2*, and *Pgc*), regulation of the host cytoskeleton (*Capn6*), and L-arginine transport (*Slc25a2*).

**Figure 3.**
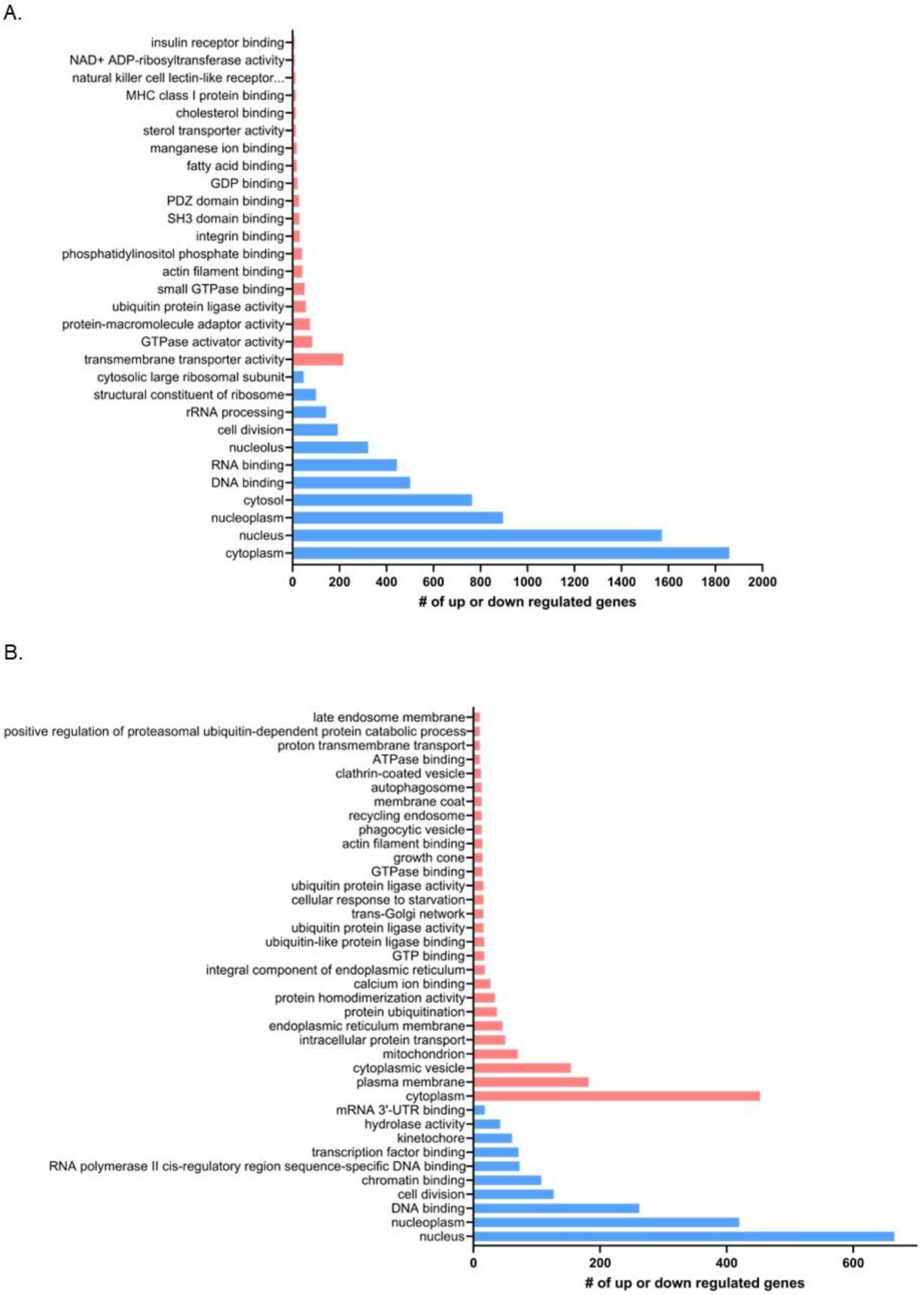
Gene ontology enrichment of differentially expressed genes. Graphs depicting both the number and directionality of differentially expressed genes categorized by gene ontology terms in the A) WT vs PBS and B) WT vs KO comparisons. GO terms with ≥ 10 genes are displayed here. (Red=downregulated; Blue=upregulated).

We then used KEGG pathway analysis to define the major pathways affected by GBS colonization in the colon. We identified 46 KEGG pathways that were significantly altered between the WT and PBS groups in the colon (false discovery rate < 0.05). Among those pathways, there were 8 involved in epithelial barrier structure and function (Figure 4A) including adherens, gap, and tight junctions, cell and focal adhesion, extracellular matrix interactions, apoptosis, and regulation of the cytoskeleton (Supplemental Figure 1A-H). There were also 14 pathways involved in immune regulation (Figure 4B) including AMPk, cAMP, chemokine, cytokine, Fc-gamma phagocytosis, Il-17, JAK-STAT, leukocyte migration, MAPk, mTOR, NFKB, NOD-like receptor, TGF-beta, and Th-17 (Supplemental Figure 2A-N). In these pathways, there are distinct gene expression differences between the WT and PBS groups, while the KO group can be considered an intermediary with some unique patterns of gene expression. Overall, our results reveal a significant effect of GBS colonization on the colonic transcriptome with major functional changes being attributed to barrier and immune function.

**Figure 4.**
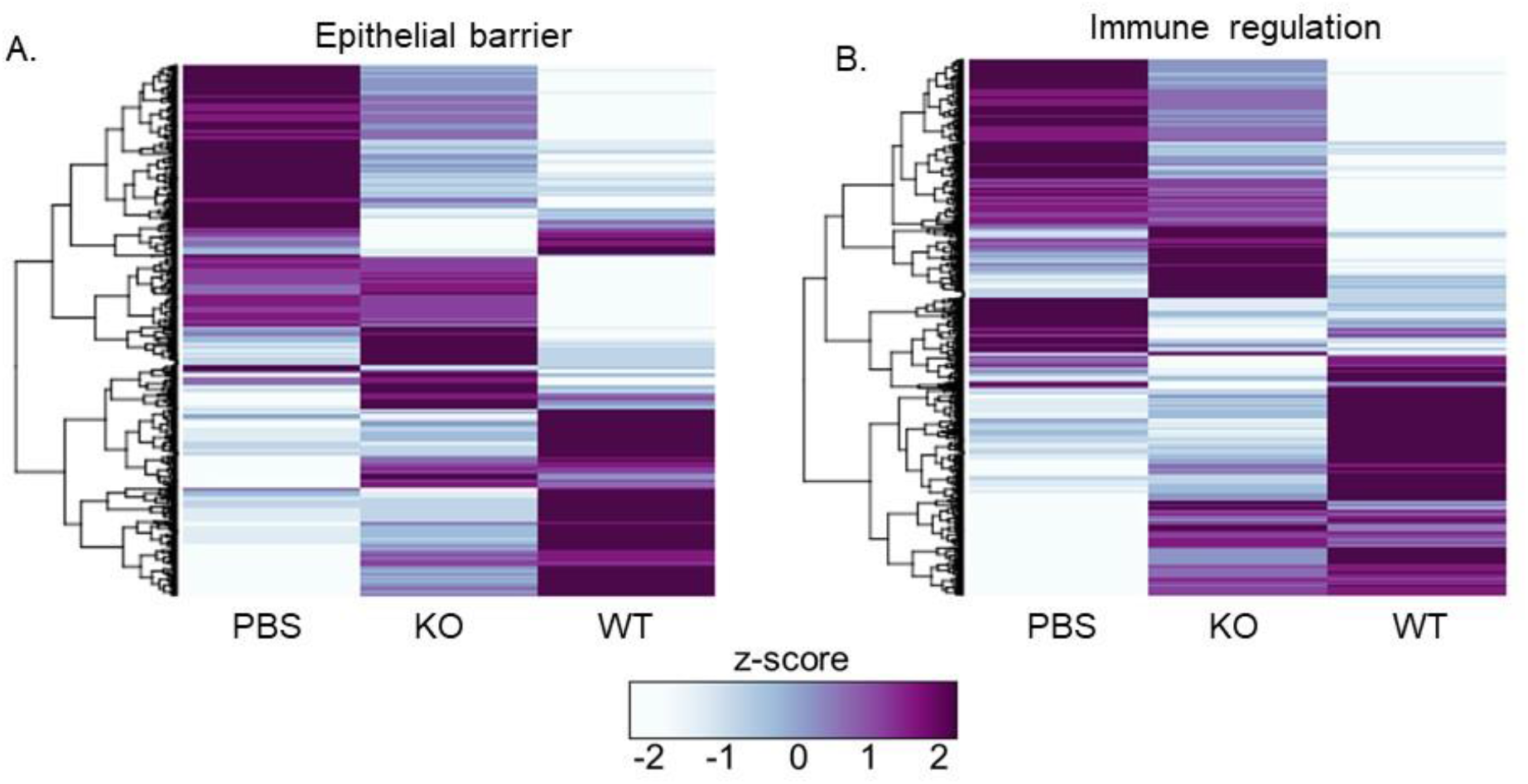
GBS alters major KEGG pathways involved in epithelial barrier function and immune regulation. Heatmaps of normalized z-scores for all genes differentially expressed in A) epithelial barrier and B) immune regulation KEGG pathways.

### βH/C is not required for intestinal colonization or GBS mortality in vivo

In our longitudinal cohort, we observed sustained GBS colonization in all animals that survived. The median time to colonization clearance was 17 days post-inoculation in the KO exposed group versus 28 days in the WT group, though this difference did not reach statistical significance (P= 0.4311, Log-rank Mantel-Cox test) (Figure 5A). Mortality occurred exclusively within the first week post-infection. We observed no statistically significant differences in mortality between WT and KO groups (P=0.4917, Log-rank Mantel-Cox test) (Figure 5B). Of note, all mortality in the KO exposed occurred in a single litter (versus in 3 of the WT exposed litters) suggesting a possible cage-related effect.

**Figure 5.**
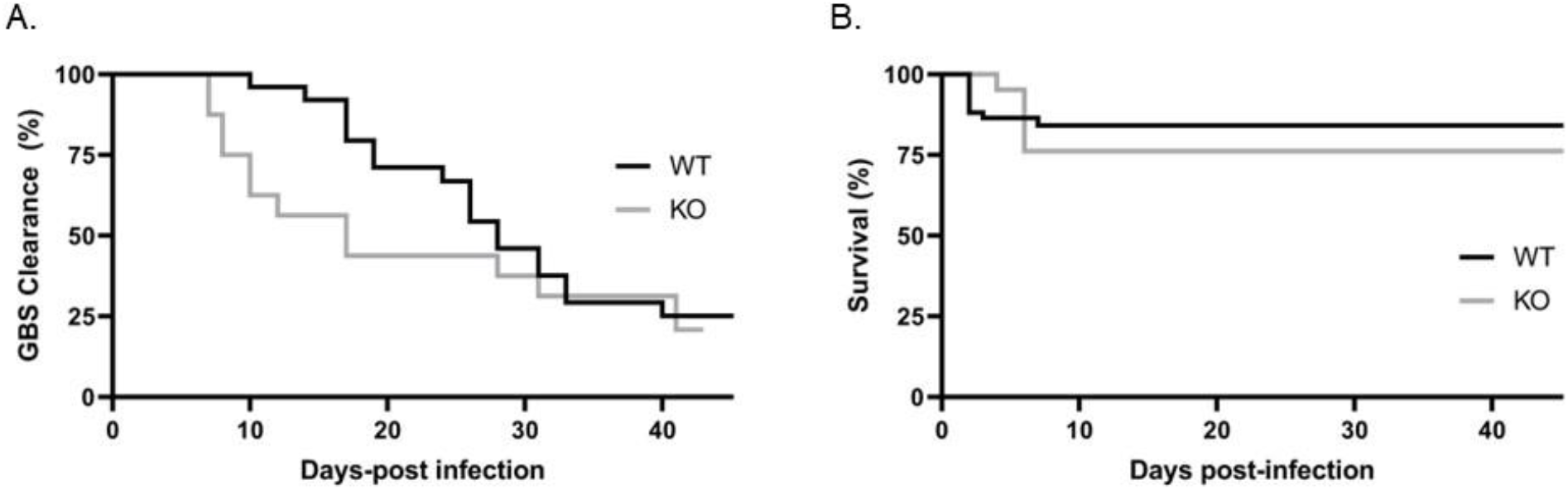
βH/C is not a major contributor to intestinal colonization or GBS mortality *in vivo*. A) Kaplan-Meier curve for mice exposed to either WT or KO (*P*=0.7431, Log-rank Mantel-Cox test). B) Kaplan-Meier curve demonstrating clearance of GBS intestinal colonization. (*P*= 0.4311, Log-rank Mantel-Cox test). (n=58 WT, n=22 KO).

## Discussion

Several enteric pathogens express toxins that compromise intestinal barrier integrity leading to subsequent dissemination and invasive disease(24, 25). Our objective was to explore GBS-host interactions in the intestinal environment and determine the specific contribution of the βH/C toxin to the development of invasive GBS disease following intestinal colonization. Our results demonstrate that expression of the toxin is not required for the establishment of GI colonization in our mouse model but can contribute to GBS disruption of critical epithelial barriers and subsequent dissemination to remote organs. We observed toxin-dependent changes in the host colonic cell transcriptome, specifically in gene pathways related to epithelial barrier function and immune regulation. These findings are consistent with previous reports indicating that GBS exploits the already compromised epithelial cell-cell junctions and underdeveloped immune repertoire in neonatal gut tissue to cross intestinal barriers (3, 5, 26, 27).

While we hypothesized that βH/C toxin contributes to GBS pathogenicity in the gut, it is noteworthy that our toxin-deficient mutant induced significant changes in the host intestinal transcriptome, established sustained gastrointestinal colonization, and was associated with mortality (although limited to a single litter) — suggesting that the toxin is the major contributor to late-onset disease. Previous investigations examining the mechanisms of GBS invasion of intestinal barriers have identified other GBS factors critical for its adhesion and translocation across intestinal epithelium including the hypervirulent GBS adhesin (HvgA)(28) and the SrtA sortase(29). The microenvironment in the infant gut also contributes to their unique vulnerability to invasive GBS disease — specifically an immature intestinal microbiome(5) and the postnatal hormonal milieu(4). The impact of these environmental factors on βH/C toxin production are unknown but a future area of exploration.

### Study limitations

Although we observed major changes in the intestinal transcriptome, we recognize that this study is limited. Validation of key changes in gene expression in human intestinal cell lines would strengthen the relevance of our findings for infants. Furthermore, examination of gene expression alone does not indicate protein production or distribution and future studies using a multi-omics approach to better understand the host pathogen dynamics are warranted.

Data reported here relating to GBS burden, dissemination, and transcriptomic changes reflect a single time point (4 days post-infection). Individual animals are likely at varying stages of disease progression (colonization, translocation, dissemination). Earlier and later time points are of interest to capture a more complete story of intestinal perturbation. Finally, determining the specific contribution of the βH/C toxin to GBS pathogenicity is challenging because it is not easily purified due to its limited solubility and instability when isolated(30). This lack of available tools (recombinant and/or stable purified toxin) is problematic and most *in vivo* studies(11–13) rely on toxin-deficient strains created via double crossover allelic exchange mutagenesis(30) such as the one used here.

## Conclusion

βH/C is a critical toxin in GBS pathogenesis, and here we show its importance in inducing transcriptomic changes in the intestinal epithelium. These results lay the foundation for further studies investigating βH/C-regulated pathways as potential targets for novel strategies to prevent GBS intestinal translocation and invasive disease in newborns.

## Acknowledgements

We would like to acknowledge funding support through the Pamela and Leslie Muma Endowed Chair in Neonatology and computational support provided by USF Omics Hub.

